# Changes in the expression of mitochondrial morphology-related genes during the differentiation of murine embryonic stem cells

**DOI:** 10.1101/644906

**Authors:** Jeong Eon Lee, Bong Jong Seo, Min Ji Han, Yean Ju Hong, Kwonho Hong, Hyuk Song, Jeong Woong Lee, Jeong Tae Do

**Affiliations:** Department of Stem Cell and Regenerative Biotechnology, KU Institute of Science and Technology, Konkuk University, 120 Neungdong-ro, Gwangjin-gu, Seoul 05029, Republic of Korea; Biotherapeutics Translational Research Center, Korea Research Institute of Bioscience and Biotechnology, Daejeon 305-806, Republic of Korea

**Author notes:** These authors contributed equally to this work (Jeong Eon Lee, Bong Jong Seo). Correspondence should be addressed to Jeong Tae Do, Ph.D.

**Keywords:** pluripotent stem cells, differentiation, mitochondria, mitochondrial morphology, mitochondrial dynamics

## Abstract

During embryonic development, cells undergo changes in gene expression, signaling pathway activation/inactivation, metabolism, and intracellular organelle structures, which are mediated by mitochondria. Mitochondria continuously switch their morphology between elongated tubular and fragmented globular via mitochondrial fusion and fission. Mitochondrial fusion is mediated by proteins encoded by *Mfn1, Mfn2*, and *Opa1*, whereas mitochondrial fission is mediated by proteins encoded by *Fis1* and *Dmn1L*. Here, we investigated the expression patterns of mitochondria-related genes during the differentiation of mouse embryonic stem cells (ESCs) in response to leukemia inhibitory factor (LIF) withdrawal. The expression of *Mfn2* and *Dnm1L* was, as expected, increased and decreased, respectively. By comparing gene expression and mitochondrial morphology, we proposed an index that could precisely represent mitochondrial changes during the differentiation of pluripotent stem cells by analyzing the expression ratios of three fusion- and two fission-related genes. Surprisingly, increased *Mfn2/Dnm1L* ratio was correlated with elongation of mitochondria during the differentiation of ESCs. Moreover, application of this index to other specialized cell types revealed that neural stems cells (NSCs) and mouse embryonic fibroblasts (MEFs) showed increased *Mfn2/Dnm1L* ratio compared to ESCs. Thus, we suggest that the *Mfn2/Dnm1L* ratio could reflect changes in mitochondrial morphology according to the extent of differentiation.

## Introduction

During embryonic development, cells undergo various changes in gene expression (Carter et al., 2003; Hamatani et al., 2004) and signaling pathways (Wang et al., 2004). Metabolism and intracellular organelle structures are also altered during development and differentiation (Folmes et al., 2012; Leese, 1995; Van Blerkom, 2004). Specifically, the organellar changes are observed during the regaining of pluripotency (also known as reprogramming) (Choi et al., 2015). For example, Folmes et al. showed that the globular shape of mitochondria progressively changed to elongated during embryonic development from zygote to somite embryo. Accordingly, metabolic features such as pyruvate oxidation, glucose oxidation, glycolysis, and the pentose phosphate pathway (PPP) were also changed dynamically (Folmes et al., 2012). These features return to the developmental early-stage status during the reprogramming process (Panopoulos et al., 2012). Some of the most dramatic changes in cells during development and differentiation occur in the mitochondria, which play essential roles in cellular processes, including energy metabolism (Seo et al., 2018), apoptosis (Suen et al., 2008), aging (Bratic and Larsson, 2013), reactive oxygen species production, calcium homeostasis, and differentiation (Wanet et al., 2015).

Mitochondria continuously change their morphology through fusion and fission in response to cellular requirements; this process is called mitochondrial dynamics or mitochondrial biogenesis (Chan, 2006; Chen and Chan, 2004; Cho et al., 2006; Westermann, 2012). In pre-implantation embryonic and pluripotent stem cells, immature mitochondria characterized by a small and globular shape with poorly developed cristae are observed (Cho et al., 2006; Chung et al., 2007; St John et al., 2005). Cells with immature mitochondria show low oxygen consumption and high levels of glycolytic enzymes (Kondoh et al., 2007). Thus, undifferentiated embryonic stem cells (ESCs) also exhibit low levels of ATP production, modest levels of antioxidant enzymes, and poor oxidant capacity (Cho et al., 2006; Chung et al., 2007; Kondoh et al., 2007). Upon differentiation of ESCs, mitochondria in these cells become elongated, showing developed cristae and dense matrices (Lonergan et al., 2007); this results in high oxygen consumption and ATP production for more efficient cellular activity (Cho et al., 2006; Chung et al., 2007; St John et al., 2005).

In mammals, mitochondrial morphology switches between elongated tubular and fragmented globular by fusion and fission, respectively (Okamoto and Shaw, 2005; Westermann, 2010). Mitochondrial fusion is mediated by the dynamin family GTPases, such as mitofusin (MFN) 1, MFN2, and optic atrophy 1 (OPA1) (Chen et al., 2003; Delettre et al., 2000; Santel and Fuller, 2001). Although the exact fusion mechanism has yet to be defined, MFN1 and MFN2 form a dimer that inserts itself into the mitochondrial outer membrane, whereas OPA1 is located in the mitochondrial inner membrane (Meeusen et al., 2006; Song et al., 2009). MFN1, MFN2, and OPA1 contain a GTPase domain, hydrophobic heptad repeat (HR) domain, and transmembrane domain (Ishihara et al., 2006; Santel and Fuller, 2001). MFN1 and MFN2 play similar roles in mitochondrial fusion and thus can functionally replace each other and form homotypic or heterotypic dimers (Chen et al., 2003; Shirihai et al., 2015).

In contrast, the major proteins related to mitochondrial fission are FIS1 (James et al., 2003; Yoon et al., 2003; Zhang and Chan, 2007) and dynamin-related protein 1 (DNM1L, also called DRP1) (Legesse-Miller et al., 2003; Otsuga et al., 1998; Smirnova et al., 1998). DNM1L is mainly located in the cytosol and recruited to the outer membrane of the mitochondria where it induces fission (Shirihai et al., 2015). FIS1 is located in the outer mitochondrial membrane and is closely related to DNM1L (James et al., 2003; Yoon et al., 2003). DNM1L can interact with other mitochondrial fission proteins, including mitochondrial fission factor (MFF) and FIS1. Interestingly, a recent study suggested that DNM1L can interact with the fusion protein MFN and facilitate MFN-mediated fusion (Shirihai et al., 2015). Although many studies have evaluated the fusion and fission of mitochondria, the mechanisms and signaling pathways that determine mitochondrial dynamics are still unclear. Activation of the mammalian target of rapamycin (mTOR) pathway by the withdrawal of LIF induced mouse ESC differentiation with suppression of pluripotent genes such as *Klf4, Oct4*, and *Nanog* (Cherepkova et al., 2016). When pluripotent stem cells differentiate, they require more energy to meet the demands of their newly acquired functions (Folmes et al., 2012)

Accordingly, mitochondria undergo dynamic remodeling during differentiation, and thus, mitochondrial morphology and metabolism are changed. Therefore, we hypothesized that expression of mitochondrial fusion- and fission-related genes may be changed toward a certain direction during the spontaneous differentiation of ESCs. Here, we quantified the expression levels of the fusion-related genes *Mfn1, Mfn2*, and *Opa1* and the fission-related genes *Fis1* and *Dnm1L* during the differentiation of murine ESCs. Here, we investigated these genes to determine whether they could be used as an index of the extent of differentiation and changes in mitochondrial morphology.

## Materials and Methods

All methods used in this study were carried out in accordance with animal care and use guidelines, and all experimental protocols were approved by the Institutional Animal Care and Use Committee of Konkuk University.

### Cell cultures

Mouse ESCs (E14tg2a) were purchased from the American Type Culture Collection (ATCC, Manassas, VA, USA) and cultured on culture dishes layered with inactivated MEFs in an ESC medium, consisting of Dulbecco’s modified Eagle’s medium (DMEM; Gibco) supplemented with 15% fetal bovine serum (FBS; HyClone), 1× Penicillin/Streptomycin/Glutamine (P/S/G; Gibco), 0.1 mM non-essential amino acids (NEAAs; Gibco), 1 mM β-mercaptoethanol (Gibco) with 1000 U/mL leukemia inhibitory factor (ESGRO, Chemicon International). MEFs were cultured in culture dishes coated with 0.15% porcine gelatin (Sigma, St. Louis, MO, USA) in MEF medium consisting of DMEM (Gibco) supplemented with 15% FBS (HyClone), 1× P/S/G (Gibco), 0.1 mM NEAAs (Gibco), and 1 mM β-mercaptoethanol (Gibco). NSCs were cultured in culture dishes coated with 0.15% porcine gelatin (Sigma) in NS medium consisting of DMEM:Nutrient Mixture F-12 (Gibco), 0.5 mg/ml bovine serum albumin (BSA; Sigma) 1% N2 supplement (Gibco), 1x NEAAs (Gibco), 1x P/S/G (Gibco), 10 ng/mL basic fibroblast growth factor (bFGF; R&D systems), and 10 ng/mL epidermal growth factor (EGF; Gibco). All cell lines were incubated at 37 °C in an atmosphere of 5% CO_2_ and maintained on tissue culture dishes (Corning, Amsterdam, The Netherlands).

### In vitro differentiation

In the pre-plating process, ESCs cultured with feeder cells were dissociated by trypsin-EDTA (0.25%) (Gibco) and transferred to a 0.15% gelatin-coated dish and incubated at 37°C in an atmosphere of 5% CO_2_ for 2 h to remove the feeder cells. Because the feeder cells attached to the gelatin-coated dish much earlier than ESCs, supernatant of the culture mostly contain ESCs without contamination of feeder cells. Supernatant was transferred another gelatin-coated dish and used as the differentiation experiment. For *in vitro* differentiation, 1 × 10^5^ ESCs were seeded in 100-mm cell culture dishes coated with 0.15% porcine gelatin (Sigma) with MEF medium. The medium was refreshed every day for 15 days of differentiation. The cells were collected by scraping for experimental analysis.

### RNA isolation and qRT-PCR

Total RNA was isolated using TRIzol reagent (Invitrogen, Carlsbad, CA, USA) according to the manufacturer’s protocol. The cDNA was then synthesized from 1 mg total RNA using SuperScript III Reverse Transcriptase (Invitrogen) and oligo(dT)20-primer (Invitrogen) according to the manufacturer’s instructions. qPCR was performed in duplicate with Power SYBR Green Master Mix (Takara, Shiga, Japan), and results were analyzed on a Roche LightCycler 5480 (Roche). Thermal cycling was carried out via 45 cycles of 10 s at 95 °C, 10 s at 60 °C, and 20 s at 72 °C. The primers for qRT-PCR are shown in Table 1.

**Table 1.**
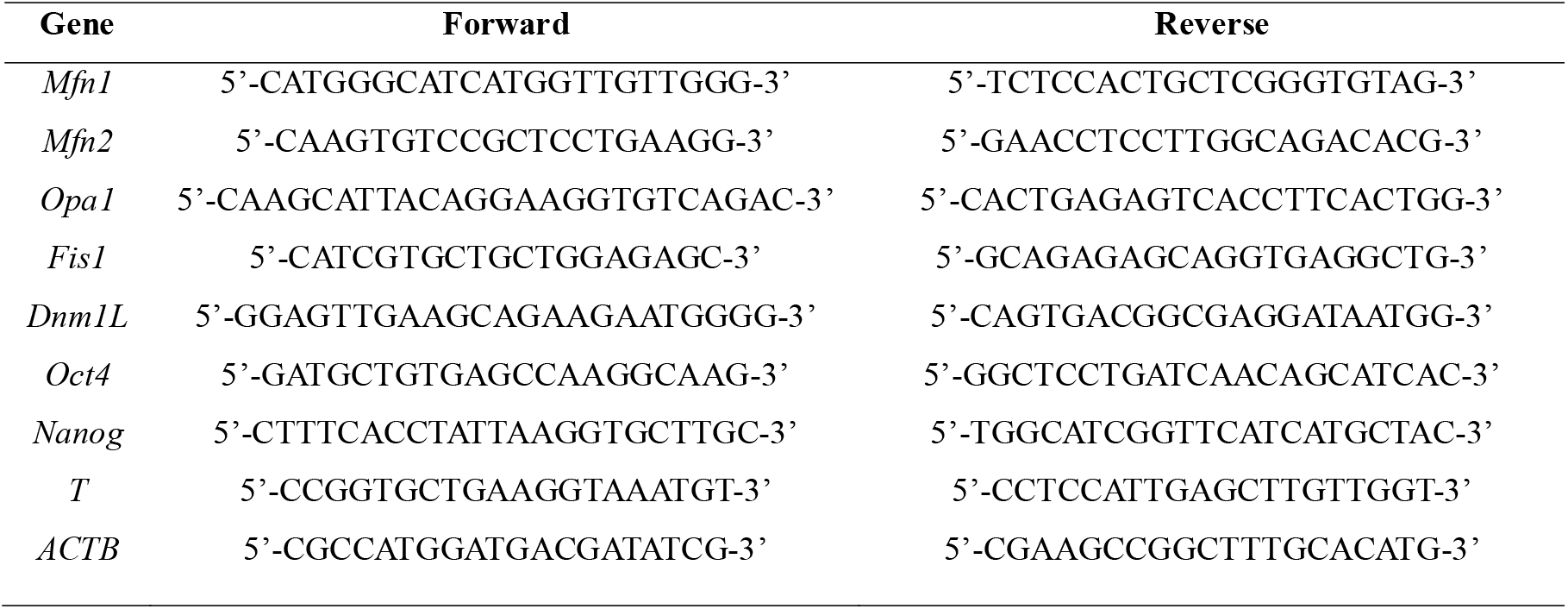
Primer sets used for quantitative RT-PCR

### Western blot analysis

Total cells were lysed using RIPA buffer (Thermo Fisher) according to the manufacturer’s instructions. Cell lysates (20 μg protein) were separated on NuPAGE 4–12% Bis-Tris Gels (Invitrogen) and transferred to polyvinyl difluoride membranes. The membranes were blocked using a blocking solution containing 5% skim milk powder and 0.05% Tween 20 in phosphate-buffered saline (PBS). The primary antibodies used in this study were as follows: anti-OCT4 (rabbit, 1:1000; Santa Cruz Biotechnology, Santa Cruz, CA, USA), anti-Nanog (rabbit, 1:1000; Abcam, Cambridge, UK), anti-MFN2 (mouse, 1:1000; Abcam), anti-DRP1 (rabbit, 1:1000; Millipore), anti-MFN1 (mouse, 1:1000; Abcam), anti-FIS1 (mouse, 1:500; Abcam), and anti-β-actin (mouse, 1:10000; Sigma). The membranes were incubated with these antibodies overnight at 4°C. Secondary antibodies were conjugated with anti-mouse IgG-peroxidase (1:10000; Sigma), anti-goat IgG-horseradish peroxidase (HRP), and anti-rabbit IgG-HRP (1:10000; Santa Cruz Biotechnology) and the membranes were incubated with these antibodies for 90 min at room temperature.

Antigens were detected using Pierce ECL Western Blotting chemiluminescent substrate (Thermo Fisher), according to the manufacturer’s instructions. Blots were then exposed to X-ray film for development and stripped for reuse of the membranes. Anti-β-actin antibody (mouse, 1:10000; Sigma) was used as a control. Densitometry of the bands for the proteins and their loading controls was performed using Image J 1.43 (NIH) software.

### Immunocytochemistry

For immunocytochemistry, cells were fixed with 4% paraformaldehyde for 20 min at room temperature. The cells were washed with PBS and then treated with PBS containing 3% bovine serum albumin and 0.03% Triton X-100 for 30 min at room temperature. The cells were then incubated with the following primary antibodies: anti-OCT4 (1:500; Santa Cruz Biotechnology), anti-Nanog (1:200; Abcam), anti-β□-tubulin (TUJ1; 1:500; R&D), anti-SMA (1:200; Abcam), anti-SOX17 (1:300; R&D), and anti-TO20 (1:200; Santa Cruz Biotechnology). Fluorescently labeled (Alexa Fluor 488 or 647; Abcam) secondary antibodies were used according to the manufacturer’s specifications. Images for anti-TOM20 staining were obtained with a confocal microscope (Zeiss).

### Electron microscopy

For transmission electron microscope (TEM) experiments, the samples were fixed in 4% paraformaldehyde (Sigma) and 2.5% glutaraldehyde (Sigma) in 0.1□M phosphate (Sigma) buffer for 24 h. After washing in 0.1□M phosphate buffer, the samples were post-fixed for 1□h in 1% osmium tetroxide (Sigma) prepared in the same buffer. The samples were dehydrated with a graded series of ethyl alcohol concentrations, embedded in Epon 812, and polymerized at 60°C for 3 days. Ultrathin sections (60–70□nm) were obtained using an ultramicrotome (Leica Ultracut UCT), collected on grids (200 mesh), and examined under a TEM (JEM 1010) operating at 60□kV, and images were recorded by a charge-coupled device camera (SC1000; Gatan).

### Mitochondrial length analysis

The images from electron microscopy were analyzed and measured by the Image J 1.43 (NIH) software for calculating the maximum (Max)/minimum/(Min) ratio of mitochondrial length. At least over fifty mitochondria were measured and analyzed per sample to obtain data.

### Statistical analysis

All experiments were performed in triplicate, and data are presented as means ± standard error of mean (SEM). Differences were assessed using unpaired or paired t-tests, and differences with p-values of less than 0.05 were considered significant.

## Results

### Changes in pluripotency- and tissue-specific markers during the differentiation of mouse ESCs

To examine changes in gene expression during the differentiation of ESCs, we randomly differentiated mouse ESCs by the withdrawal of LIF without feeder cells for 0, 3, 6, 9, 12, and 15 days (Fig. 1a). Dome-like colonies of undifferentiated ESCs became flat and showed changes in morphology. First, we checked whether ESCs were properly differentiated *in vitro* after 15 days in differentiation medium. Immunocytochemistry analysis showed that OCT4 and NANOG, which were expressed in undifferentiated ESCs, were silenced at day 15 after differentiation. Differentiation markers, neuron-specific class III β-tubulin (TUJ1), smooth muscle actin (SMA), and SRY-box 17 (SOX17) for ectodermal, mesodermal, and endodermal cells, respectively, were not detectable in undifferentiated ESCs at day 0, but were detected after differentiation at day 15, indicating that ESCs lost pluripotency and were differentiated into various cell types, including all three germ layers (Fig. 1b).

**Fig. 1.**
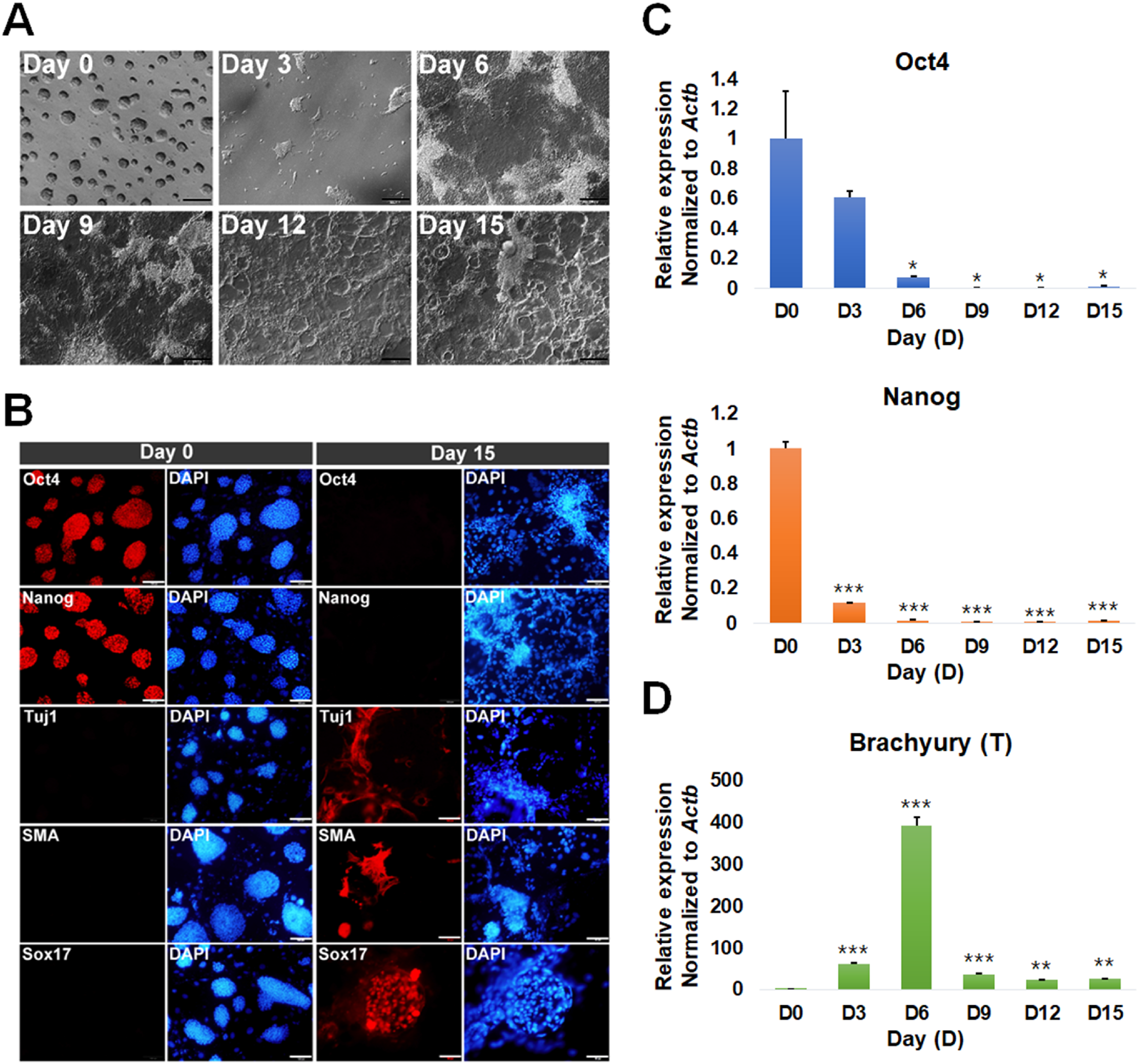
*In vitro* differentiation and characterization of ESCs. (a) Phase-contrast images of ESCs and differentiating cells on days 0, 3, 6, 9, 12, and 15. Scale bars = 200 μm. (b) Immunofluorescence images of the pluripotency markers OCT4 and NANOG and differentiation markers for ectodermal (TUJ1), mesodermal (SMA), and endodermal (SOX17) cells in undifferentiated ESCs and ESC-derived differentiated cells on days 0 and 15. Nuclear was counterstained with DAPI. Scale bars = 100 μm. (c-d) Quantitative RT-PCR analysis of ESCs and differentiating cells on days 0, 3, 6, 9, 12, and 15 (D0, D3, D6, D9, D12, and D15). Data are presented as mean ± SEM for n = 3 independent experiments. (c) Pluripotent marker *Oct4* and *Nanog* expression on day 0 to 15 (d) Differentiation marker *T* expression from day 0 to 15. t-test: *p < 0.05, **p < 0.01, and ***p < 0.001.

Next, we evaluated the expression levels of pluripotency and differentiation markers by quantitative reverse transcription polymerase chain reaction (qRT-PCR) analysis (Fig. 1c, d). The expression levels of the pluripotency markers *Oct4* and *Nanog* were gradually decreased as ESCs were differentiated (Fig. 1c). In contrast, early mesoderm marker *T* (also known as Brachyury) expression was increased until day 6 post-differentiation and then downregulated afterward (Fig. 1d), indicating that early mesoderm cells appeared at day 6 after differentiation and then differentiated further. Taken together, our findings demonstrated that ESCs differentiated gradually and lost pluripotency over 15 days upon LIF withdrawal from the ESC medium.

### Changes in mitochondrial morphology during the differentiation of mouse ESCs

Mitochondrial morphology was expected to change from fragmented to elongated shapes during the differentiation of ESCs. Thus, we examined mitochondrial morphology by immunostaining using antibodies targeting translocase of outer membrane 20 (TOM20), which is in the outer mitochondrial membrane. In ESCs, small mitochondria (green dots in Fig. 2a) were evenly distributed in the cytoplasm, indicating that the fragmented form of mitochondria was the major form in ESCs. However, green dots became larger beginning on day 6 after differentiation of ESCs (Fig. 2a). This observation was also accurately confirmed by electron microscopic analysis (Fig. 2b). Undifferentiated ESCs had primarily globular mitochondria with immature cristae, and this morphology was maintained until day 3 of differentiation. From day 6 to 15, mitochondria were gradually elongated and showed mature cristae (Fig. 2b). We measured the Max and Min axes of mitochondria to quantify mitochondrial length (Fig. 2c-d). The Max/Min ratios of mitochondria was 1.51, 1.66, 3.65, 3.92, 4.30, and 6.87 on days 0, 3, 9, 12, and 15, respectively (Fig. 2e). These data clearly demonstrated that the globular and immature mitochondria in ESCs became elongated and showed developed mature cristae after the spontaneous differentiation.

**Fig. 2.**
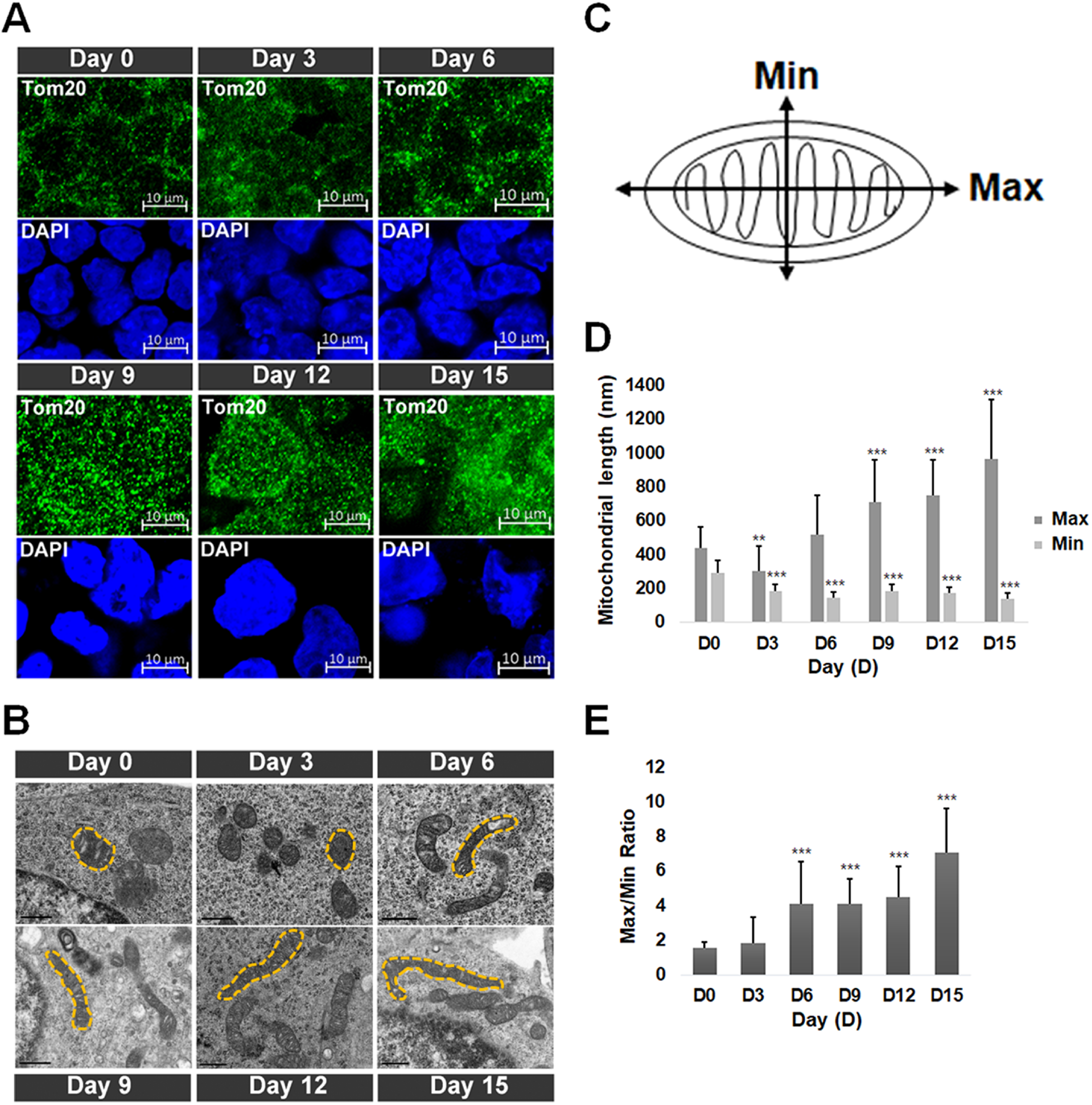
Changes in mitochondrial morphology during the differentiation of ESCs. (a) Analysis of mitochondrial morphology by staining of mitochondria using anti-TOM20 antibodies on days 0, 3, 6, 9, 12, and 15 after differentiation of ESCs. Green dots represent mitochondria. Nuclear was counterstained with DAPI. Scale bars = 10 μm. (b) Electron microscopic images of mitochondria in undifferentiated ESCs and differentiated cells from ESCs on days 0, 3, 6, 9, 12 and 15. Yellow dotted lines represent mitochondrial morphologies. Scale bars = 0.5 μm. (c) Measurement of the maximum (Max) and minimum (Min) axes of mitochondria. (d) The mitochondrial length (nm) in ESCs on differentiating days 0, 3, 6, 9, 12 and 15. (e) The Max/Min ratios in ESCs and differentiating cells on days 0, 3, 6, 9, 12, and 15. Data are presented as mean ± SEM for n = 30 independent experiments. t-test: *p < 0.05, **p < 0.01, and ***p < 0.001.

### Changes in mitochondrial morphology-related genes during the differentiation of mouse ESCs

Because mitochondria were elongated during the differentiation of ESCs, we predicted that fusion-related genes would be up-regulated, and fission-related genes would be down-regulated during this differentiation. Thus, we examined mitochondrial morphology-related genes by qRT-PCR analysis. Unexpectedly, the fusion-related gene *Mfn1* was progressively downregulated, whereas *Mfn2* and *Opa1* expression patterns fluctuated at day 3 post-differentiation (Fig. 3a). Next, we evaluated the expression of two fission-related genes, *Dnm1L* and *Fis1* (Fig. 3b); consistent with the reduced mitochondrial fission, which resulted in enhanced mitochondrial elongation, the expression of Dnm1L decreased as mouse ESCs underwent spontaneous differentiation (Fig. 3b). However, *Fis1* expression did not decrease but rather increased after slight downregulation at the beginning of differentiation (Fig. 3b). This result supported previous findings that FIS1 is less related to mitochondrial fission than DNM1L and functions to recruit DNM1L to the mitochondria (Loson et al., 2013; Zhang et al., 2016). Collectively, our results showed that *Mfn2* and *Dmn1L* mRNA expression levels reflected mitochondrial elongation with ESC differentiation.

**Fig. 3.**
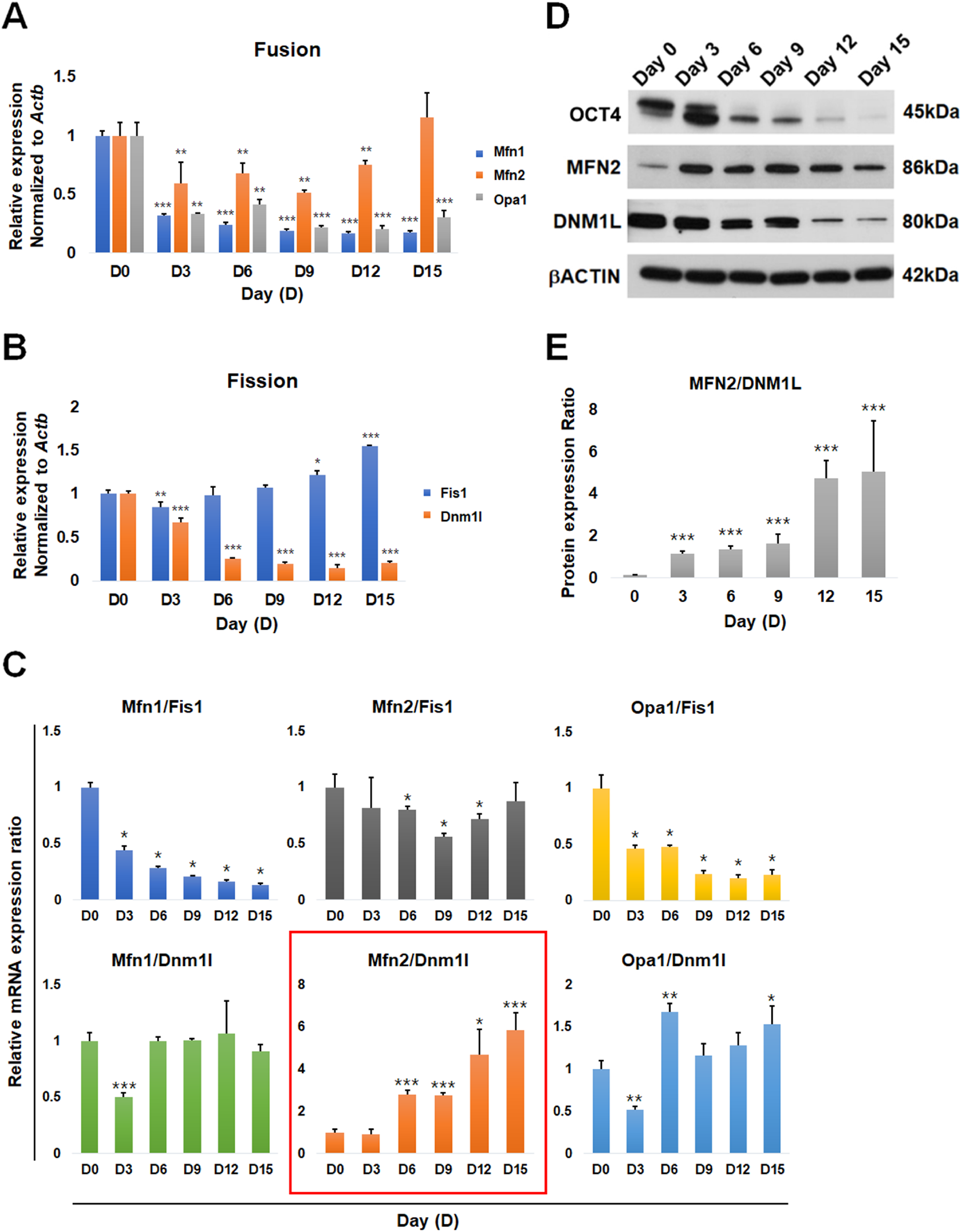
Changes in the expression levels of mitochondrial fusion- and fission-related genes and in indexes from combinations of fusion- and fission-related genes. Quantitative RT-PCR analysis of (a) mitochondrial fusion-related genes (*Mfn1, Mfn2*, and *Opa1*) and (b) mitochondrial fission-related genes (*Fis1* and *Dnm1L*) on days 0, 3, 6, 9, 12, and 15 after differentiation of ESCs. Gene expression levels were normalized to the expression of *Actb*. (c) Six combinatorial ratios from the analysis of gene expression levels between three fusion- and two fission genes. (d) Western blotting for the expression of OCT4, MFN2, and DNM1L. β-actin was used as a control for other proteins. (e) MFN2/DNM1L protein ratios were gradually increased according to the elapsed time after ESC differentiation. All data are presented as mean ± SEM for n = 3 independent experiments. t-test: *p < 0.05, **p < 0.01, and ***p < 0.001.

Accordingly, we next investigated the levels of MFN2 and DNM1L proteins by western blotting in ESCs (day 0) and in differentiated cells on days 3, 6, 9, 12, and 15 (Fig. 3d). As a control for ESC spontaneous differentiation, the expression of the pluripotency marker OCT4 gradually decreased during differentiation and was almost undetectable at day 15. MFN2 was expressed at a low level in ESCs but showed a dramatic increase in expression upon spontaneous differentiation. Furthermore, MFN2 protein levels showed a fluctuating expression pattern during differentiation, similar to the qRT-PCR data (Fig. 3a and 3d). DNM1L protein levels decreased gradually during the differentiation of ESCs, similar to the result of qRT-PCR analysis (Fig. 3b and 3d). Thus, we observed similar changes in the mRNA and protein levels for these two targets.

### Establishment of indexes representing the extent of differentiation

Given that many genes involved in mitochondrial fusion and fission are not correlated with mitochondrial morphology, we next attempted to find an index to represent mitochondrial morphology. Based on the qRT-PCR data, we analyzed the ratios between three fusion- and two fission-related genes during the differentiation of ESCs. A total of 6 combinatorial ratios for three fusion- and two fission-related genes were analyzed (Fig. 3c). One of these, the *Mfn2/Dnm1l* ratio, was interesting because it was similar to the mitochondrial length Max/Min ratio during the differentiation of ESCs (Fig. 2d and 3c in red square). Next, we confirmed the protein expression level in the *MFN2/DNM1L* ratio (Fig. 3e). Indeed, the MFN2/DNM1L ratio was also similar to the mRNA expression and mitochondrial length Max/Min ratio. Thus, we concluded that the mitochondrial changes during ESC spontaneous differentiation could be reflected by the progressive increase in the *Mfn2/Dnm1l* ratio.

### Mfn2/Dnm1L index represented mitochondrial morphology according to the extent of differentiation

To investigate whether this index was applicable to other cell types, we compared this ratio among ESCs, neural stem cells (NSCs) and mouse embryonic fibroblasts (MEFs). We chose these cell types because ESCs are undifferentiated, NSCs are less specialized, and MEFs are differentiated. Only ESCs express high levels of *Oct4* (Fig. 4b). Although the mRNA expression level of the mitochondrial fusion-related gene *Mfn2* was slightly changed in these cell types, the mRNA expression level of the mitochondrial fission-related gene *Dnm1l* was gradually decreased in NSCs and MEFs (Fig. 4c, 4d), consistent with the observed changes in mitochondrial morphology (Fig. 4a). Next, we applied the *Mfn2/Dnm1l* ratio to these cell types, with adjustment to a ratio of 1.0 for ESCs. Interestingly, the adjusted *Mfn2/Dnm1L* ratios were 1.0, 2.97, and 3.86 in ESCs, NSCs, and MEFs, respectively. Next, we also confirmed MFN2 and DNM1L protein expression (Fig. 4f). As expected, the DNM1L protein expression levels were decreased in NSCs and MEFs, compared with those in ESCs. Interestingly, the MFN2 protein expression levels were gradually increased in NSCs and MEFs. The *MFN2/DNM1L* protein expression ratios were 0.16, 2.94, and 4.61 in ESCs, NSCs, and MEFs, respectively. Thus, we concluded that the *Mfn2/Dnm1l* ratio at the mRNA and protein expression levels corresponded to the length of mitochondria (Fig. 4e-g).

**Fig. 4.**
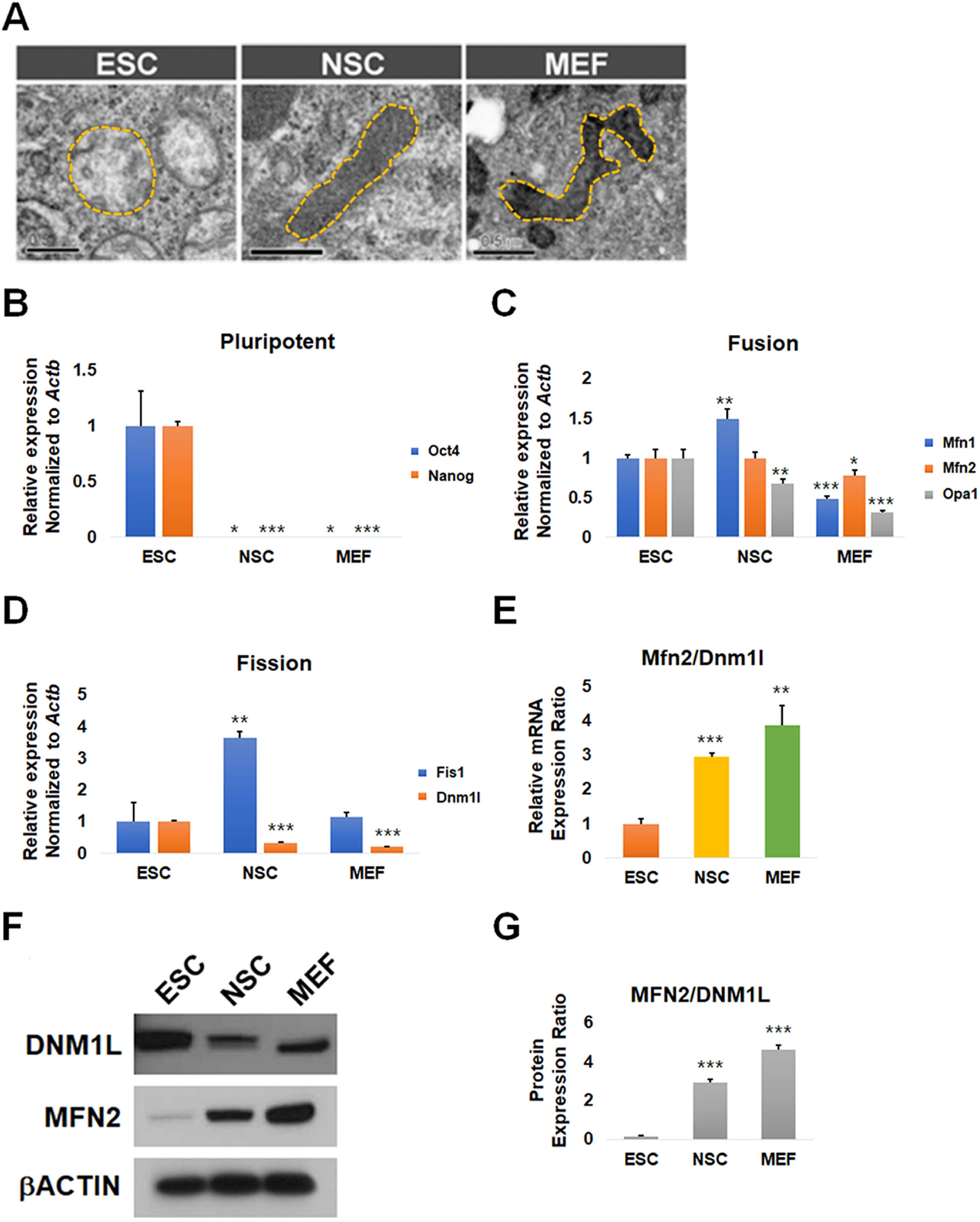
*Mfn2/DnmlL* ratios in ESCs, NSCs, and MEFs. (a) Electron microscopic images of mitochondria in undifferentiated ESCs, NSCs, and MEFs. Yellow dotted lines represent mitochondrial morphologies. Scale bars = 0.5 μm. Quantitative RT-PCR analysis of (b) pluripotency-, (c) mitochondrial fusion-, and (d) mitochondrial fission-related genes in ESCs, NSCs, and MEFs. Gene expression levels were normalized to the expression of *Actb*. (e) *Mfn2/Dnm1L* gene expression ratios in ESCs, NSCs, and MEFs. (f) Western blotting for the expression of DNM1L and MFN2 in ESCs, NSCs, and MEFs. β-actin was used as a control for other proteins. (g) MFN2/DNM1L protein ratios in ESCs, NSCs, and MEFs. All data are presented as mean ± SEM for n = 3 independent experiments. t-test: *p < 0.05, **p < 0.01, and ***p < 0.001.

## Discussion

In this study, we investigated the dynamics of mitochondrial morphology-related genes, which are responsible for mitochondrial fusion and fission, during the differentiation of mouse ESCs. Given that differentiated cells contain elongated mitochondria and ESCs contain globular mitochondria (Seo et al., 2018), we expected to observe increases in the expression of mitochondrial fusion-related genes (*Mfn1, Mfn2*, and *Opa1*) and decreases in the expression of fission-related genes (*Fis1* and *Dnm1L*).

This is because several reports have shown that the overexpression of fusion-related genes such as *Mfn1/Mfn2* (Chen et al., 2003) and *Opa1* (Frezza et al., 2007) induces mitochondrial elongation in MEFs. In this context, the overexpression of fission-related genes such as *Drp1* (Yamamori et al., 2015) and *Fis1* (Otera et al., 2010) induces mitochondrial fragmentation in MEFs and HeLa cells, respectively. However, there were only minor changes in the expression levels of *Mfn1* and *Opa1* during differentiation, and changes in the expression of *Fis1* were opposite to the expected results. Actually, FIS1 has been reported to have a weaker effect on mitochondrial fission than DNM1L in MEFs but not in HeLa (Loson et al., 2013; Otera et al., 2010; Yamamori et al., 2015) possibly due to the different mechanisms of mitochondrial morphology regulation in humans and mice. Fig. 3a shows that the expression pattern of *Mfn2* better reflects mitochondrial morphology than that of *Mfn1* or *Opa1* during ESC spontaneous differentiation. A loss-of-function experiment in a previous study showed that *Mfn1* or *Mfn2* deficiency results in fragmented mitochondrial morphology and loss of *Mfn2* has more dramatic effects than loss of *Mfn1* (Chen et al., 2003). Moreover, OPA1, processed for mitochondrial fusion by protease isoenzymes and OMA1, might not reflect the morphology of mitochondria (Ehses et al., 2009). This is because various proteins affect the processing of OPA1 protein for its function in the mitochondrial fusion. Only *Mfn2* and *Dnm1L* expression patterns were, as expected, increased and decreased, respectively.

Next, we aimed to identify a special index that gradually increased during ESC spontaneous differentiation. We found that a gradual increase in the *Mfn2/Dnm1L* ratio was closely related to mitochondrial elongation during the elapsed time after the differentiation of ESCs. Furthermore, we found that this index could be applied to other cell types, including NSCs and MEFs. More specialized cells showed higher *Mfn2/Dnm1L* ratios than ESCs.

There were some limitations to our research. Firstly, technologies such as High-Content Imaging and Machine Learning were recently developed for the analysis of the shapes of mitochondria (Leonard et al., 2015). This technical method allows the analysis of the mitochondria by further subdividing the shape of the mitochondria e.g., fragmented, rods, networks, and large/round. However, the functions of the genes that affect the dynamic of mitochondria have yet to be identified and therefore cannot explain how mitochondria form a network (Seo et al., 2018). Therefore, we focused on classifying the mitochondria in just two categories, i.e., fragmented and elongated. Second, some cell types at final stages of differentiation such as erythropoietic cells (Jensen et al., 2019; Liu et al., 2017) and hepatocytes (Das et al., 2012) have fragmented mitochondria. If ESCs were differentiated by LIF-withdrawal, various cell types will be observed among differentiated cell population (Cherepkova et al., 2016; Duval et al., 2000), including erythropoietic cells and hepatocytes. The fragmentation of mitochondria related to the specific role of cells cannot be limited to gene expression, which controls the dynamic of mitochondria. Thus, the tendency of gene expression patterns can only be interpreted as the indexes that predict the degree of differentiation and shape of mitochondria, because of the random differentiation process (i.e., no lineage-specific differentiation). Thus, our findings suggested that this ratio could also be used as an index for mitochondrial morphology during differentiation.

## Acknowledgments

This research was supported by the Basic Science Research Program through the National Research Foundation of Korea (NRF) funded by the Ministry of Science, ICT and Future Planning of the Republic of Korea (grant nos. 2016M3A9B6946835 and 2015R15A1009701).

## Author contribution statement

JEL, KH, BJS, and JTD wrote the main manuscript text and designed the study. JEL, BJS, and YJH, MJH performed experiments and analyzed the data. JEL, BJS, HS, and JWL performed data analysis. All authors reviewed the manuscript.

